# Circulating small non-coding RNAs associated with age, sex, smoking, body mass and physical activity

**DOI:** 10.1101/247155

**Authors:** Trine B Rounge, Sinan U Umu, Andreas Keller, Eckart Meese, Giske Ursin, Steinar Tretli, Robert Lyle, Hilde Langseth

**Affiliations:** Department of Research, Cancer Registry of Norway, Oslo, Norway.; Department of Clinical Bioinformatics, Saarland University, 66041, Saarbruecken, Germany.; Department of Human Genetics, Saarland University, 66421, Homburg/Saar, Germany.; Department of Medical Genetics, Oslo University Hospital and University of Oslo, Oslo, Norway; PharmaTox Strategic Research Initiative, School of Pharmacy, University of Oslo, Oslo, Norway

**Keywords:** small non-coding RNA, serum, age, sex, storage time, smoking, BMI, body mass, physical activity, RNA-sequencing

## Abstract

Non-coding RNAs (ncRNA) are regulators of cell functions and circulating ncRNAs from the majority of RNA classes, such as miRNA, tRNA, piRNAs, lncRNA, snoRNA, snRNA and miscRNAs, are potential non-invasive biomarkers. Understanding how non-disease traits influence ncRNA expression is essential for assessing their biomarker potential.

We studied associations of common traits (sex, age, smoking, body mass, physical activity, and technical factors such as sample storage and processing) with serum ncRNAs. We used RNAseq data from 526 donors from the Janus Serum Bank and traits from health examination surveys. We identified associations between all RNA classes and traits. Ageing showed the strongest association with ncRNA expression, both in terms of statistical significance and number of RNAs, regardless of RNA class. Serum processing modifications and storage times significantly altered expression levels of a number of ncRNAs. Interestingly, smoking cessation generally restored RNA expression to non-smoking levels, although for some isomiRs, mRNA fragments and tRNAs smoking-related expression levels persisted.

Our results show that common traits influence circulating ncRNA expression. Therefore it is clear that ncRNA biomarker analyses should be adjusted for age and sex. In addition, for specific ncRNAs identified in our study, analyses should also be adjusted for body mass, smoking, physical activity and serum processing and storage.

## Introduction

Approximately two thirds of the mammalian genome is transcribed to produce different RNA classes, the majority of which are non-coding RNAs (ncRNA)^1,2^. The major ncRNA classes are microRNA (miRNA), transfer RNA (tRNA), piRNAs, long non-coding RNA (lncRNA), small nucleolar (snoRNA), small nuclear RNA (snRNA) and miscellaneous RNA (miscRNAs). In addition, fragments and isoforms of RNAs may have important biological roles independent of the canonical, full-length RNAs from which they derive^3–5^. Circulating small non-coding RNAs (sncRNA) are secreted from cells, either bound to RNA binding proteins^6^, high-density lipoproteins^7^, within extracellular vesicles or released during cell death^8^. sncRNAs are protected from degradation, and miRNAs, the most studied sncRNA class, have been identified in all body fluids^9–11^. Aberrant expression of small and long regulatory non-coding RNAs are related to many diseases^12,13^.

Circulating ncRNAs have considerable potential as minimally invasive cancer biomarkers^14–19^. However, few if any have reached their translational potential. To be reliably used as biomarkers, variation and traits that influence sncRNA expression levels need to be identified in non-diseased individuals. Common traits may include age, sex, smoking, body mass and physical activity. Technical factors, such as sample processing and storage, may also influence RNA levels^20,21^. Almost all studies to date have focused on miRNAs, and have inadequate sample sizes to assess normal variation and identify the effects of traits on expression.

sncRNAs may be encoded on the sex chromosomes^22^ and sex-specific miRNA expression patterns have been shown in tissues^23^. Several steroid sex hormones, such as estradiol, progesterone and testosterone have been found to directly or indirectly regulate miRNA expression^24–26^ or Argonaute, Drosha and Dicer, the major enzymes of miRNA biogenesis^27^. Some isomiRs have also been shown to be sex-specific^28^.

Ageing is more strongly associated with circulating miRNA expression than sex. The miRNAs significantly influenced by age included hsa-miR-1284, hsa-miR-93–3p, hsa-miR-1262, hsa-miR-34a-5p, and hsa-miR-145–5p^29^. This is in agreement with the first observations of altered circulating miRNA levels during ageing showing an increase in miR-34a in the plasma of old mice^30^. 127 of 150 miRNAs analysed were shown to be affected by age in a study on whole-blood from 5221 individuals. A miRNA age prediction model was developed using this large dataset and the miRNA predicted age correlated with chronological age with an r=0.61 adjusted for cell type composition^31^. Transforming growth factor beta signalling has been suggested as one of the main pathways regulated by the differentially expressed circulating miRNAs^32^. However, cellular senescence, ageing and age-related diseases, have been associated with alterations in miRNA expression that could have multiple physiological effects. Whether the changes have an etiological origin or are a consequence of deleterious age-induced dysfunctions is still unknown^33^.

A large study (N=226) showed that smoking alters circulating miRNA expression. There was no significant finding when comparing former to never smokers^34^. A study of small airway epithelium from 10 smokers identified differences in miRNA expression after smoking cessation which persisted in 8 out of the 34 (FDR<0.05) smoking-related miRNAs, with the Wnt/β-catenin signalling pathway being the most significant pathway^35^. A smaller study of 12 never-smokers and 28 smokers, all males, identified 35 differentially expressed miRNAs, and target enrichment analyses identified the immune system and hormone regulation as possible pathways differing between the groups^36^.

Studies of differential miRNA expression related to body mass have mostly focused on adipose tissue, and a small number of miRNAs were found to be differentially expressed in individuals with obesity and type 2 diabetes mellitus (T2DM)^37^. These miRNAs influence the expression and secretion of inflammatory proteins. Ameling *et al*. found 19 of 179 miRNAs to be associated with body mass in 372 population-based samples, 12 miRNAs were age-associated and 7 were sex-associated^38^.

Physical activity-related miRNAs have mainly been found in intervention studies, identifying 4 to 23 differentially expressed miRNAs^39–41^. Oppose to changes in circulating miRNAs in acute exercise, the changes of circulating miRNAs in chronic exercise remain unclear^42^. A positive linear correlation between training-induced changes in circulating miR-20a levels and changes in VO2max has been shown, suggesting potential biomarkers of cardiorespiratory fitness trainability^43^.

With few exceptions, miRNAs are the only ncRNA class that have been studied in relationship to sex, age, body mass and physical activity. In addition, small sample sizes in most of these studies hampers discovery, and the widespread use of disease-related samples may introduce bias. tRNA, piRNAs, lncRNA, snoRNA, snRNA, miscRNAs and their isoforms may be potential biomarkers as long as they are stable, quantifiable, and population variation due to common traits is known.

In this study, we explore the relationship between sex, age, smoking, body mass, physical activity, technical factors and circulating ncRNA expression levels. We use RNAseq to high depth (on average 18 mill. sequences) from a large serum sample set (N=526) of cancer-free donors from the Janus Serum Bank (JSB)^44,^. This data, combined with high-quality survey information^45^, provides a unique opportunity to identify trait associations that might influence sncRNA biomarker potential.

## Results

### Significant trait associations

We produced RNAseq expression profiles for sncRNAs 17 to 47 nucleotides long in serum samples from cancer-free JSB donors and analysed associations with age, sex, body mass, smoking, physical activity or technical factors (blood donor group, see Material and Method section). We analysed 27251 sncRNAs, including 15217 mRNA fragments. 2127 trait associations were significant using an adjusted p-value<0.05 cut-off (Table 1 and Supplementary Table S1). When applying a stricter cut-off (adjusted p-value<0.001), 651 of the sncRNAs showed trait associations. Age had the highest number of trait associations with 1340 sncRNAs (adjusted p-value<0.05). With a stricter cut-off (adjusted p-value<0.001), 554 sncRNAs were associated with age (Supplementary Information Table S2 and S3). Only three sncRNAs were significantly associated with blood donor group (adjusted p-value<0.001; Table 1). We adjusted for age in the analyses of sex, body mass, smoking and physical activity. Age-adjustment increased the significant associations from 33 to 439 for sex, 44 to 411 for body mass, 5 to 208 for physical activity and 11 to 182 for smoking (adjusted p-value<0.001; Table 1 and Supplementary Table S4). In total, 1240 sncRNAs were associated with sex, body mass, physical activity or smoking after age adjustment (p-values < 0.001; Supplementary Table S5).

**Table 1:** Number of differentially expressed small non-coding RNAs (adjusted p-value <0.001) The number of significantly (p-value adjusted for multiple testing <0.001) differential expressed sncRNAs associated with age, body mass, blood donor group, sex, physical activity and smoking. Age significantly influenced expression, thus the number of RNA molecules associated with body mass, physical activity and smoking were also presented with age adjustment.

| RNA | age | sex | Sex<br>+ age | body<br>mass | body mass<br>+ age | physical<br>activity | physical<br>activity +<br>age | smoking | smoking+a<br>ge |
| --- | --- | --- | --- | --- | --- | --- | --- | --- | --- |
| miRNAs | 11 | 0 | 11 | 0 | 0 | 0 | 0 | 1 | 2 |
| isomiRs | 26 | 4 | 21 | 7 | 26 | 0 | 17 | 2 | 18 |
| tRNAs | 3 | 2 | 6 | 0 | 27 | 1 | 5 | 0 | 3 |
| tRNA<br>fragments | 1 | 0 | 27 | 0 | 28 | 0 | 10 | 0 | 0 |
| piRNAs | 6 | 1 | 59 | 3 | 24 | 0 | 7 | 0 | 3 |
| lncRNAs | 30 | 3 | 53 | 5 | 43 | 0 | 24 | 0 | 31 |
| snoRNAs | 1 | 1 | 4 | 1 | 1 | 0 | 0 | 0 | 5 |
| snRNAs | 48 | 0 | 1 | 5 | 5 | 0 | 1 | 8 | 12 |
| mRNAs | 428 | 22 | 257 | 23 | 257 | 4 | 144 | 0 | 108 |
| **Total** | **554** | **33** | **439** | **44** | **411** | **5** | **208** | **11** | **182** |

Hierarchical clustering of adjusted p-values for all associations were visualized using heatmaps (Figure 2). The age-associations were more numerous and with higher −log p-values than other traits and are an outgroup in the vertical dendrogram. Associations with sex, body mass, smoking and physical activity with age as a covariate, showed more and stronger associations. Notably, piRNAs were associated with sex, after adjusting for age (Supplementary Figure S1).

**Figure 1:**
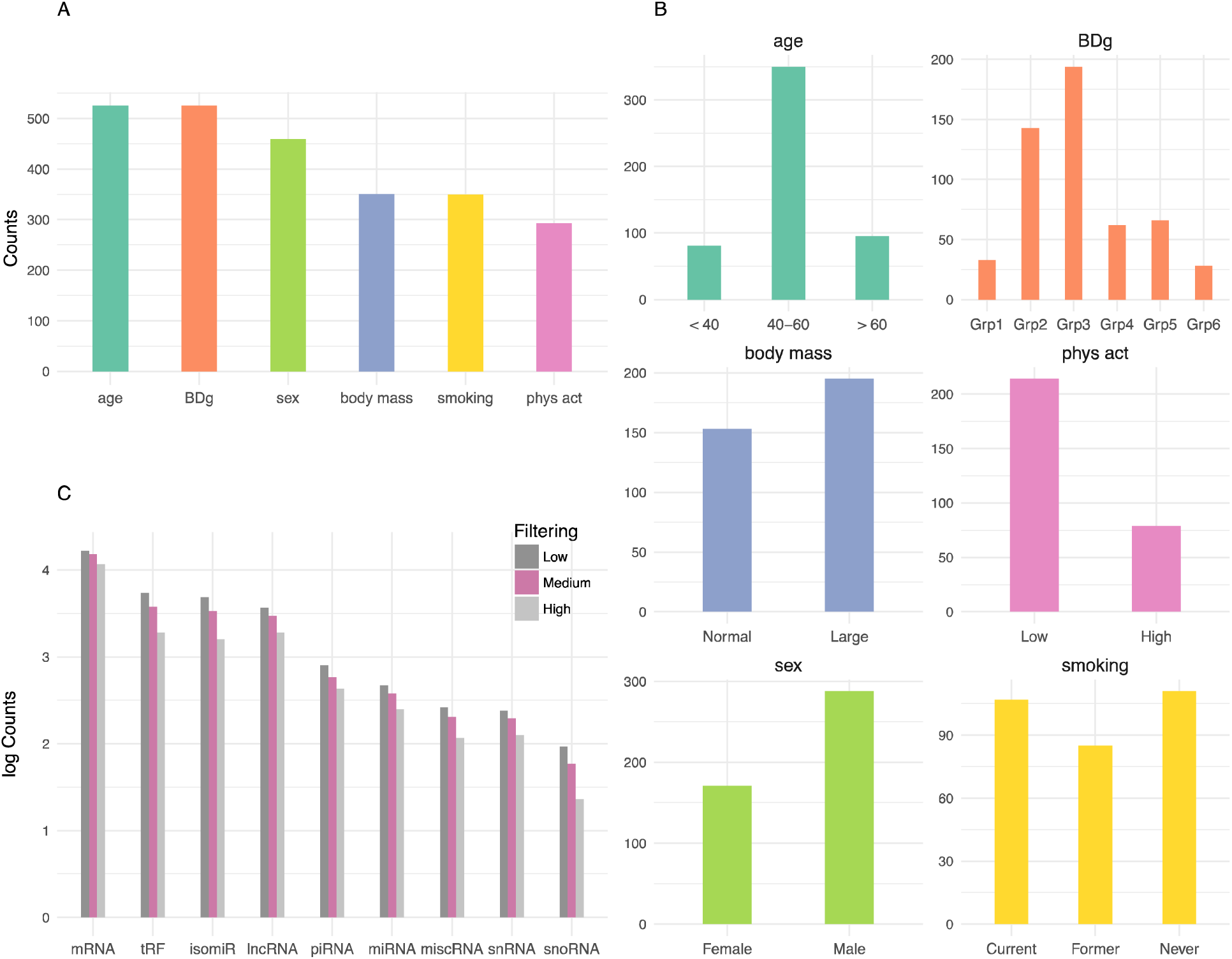
Associations between traits and small non-coding RNAs (sncRNA) were investigated in samples from in total 526 Janus Serum Bank donors, including 156 Red Cross Blood Donors and 370 Health Examination Blood Donors. A) The total number of samples included in each trait analyses after excluding samples with missing data and low sncRNA yielding samples. B) Number of samples in each category for age, blood donor group (BDg), body mass, physical activity (phys act), sex and smoking. C) Number of sncRNA counts on log scale after less stringent, medium and high stringent filtering.

**Figure 2:**
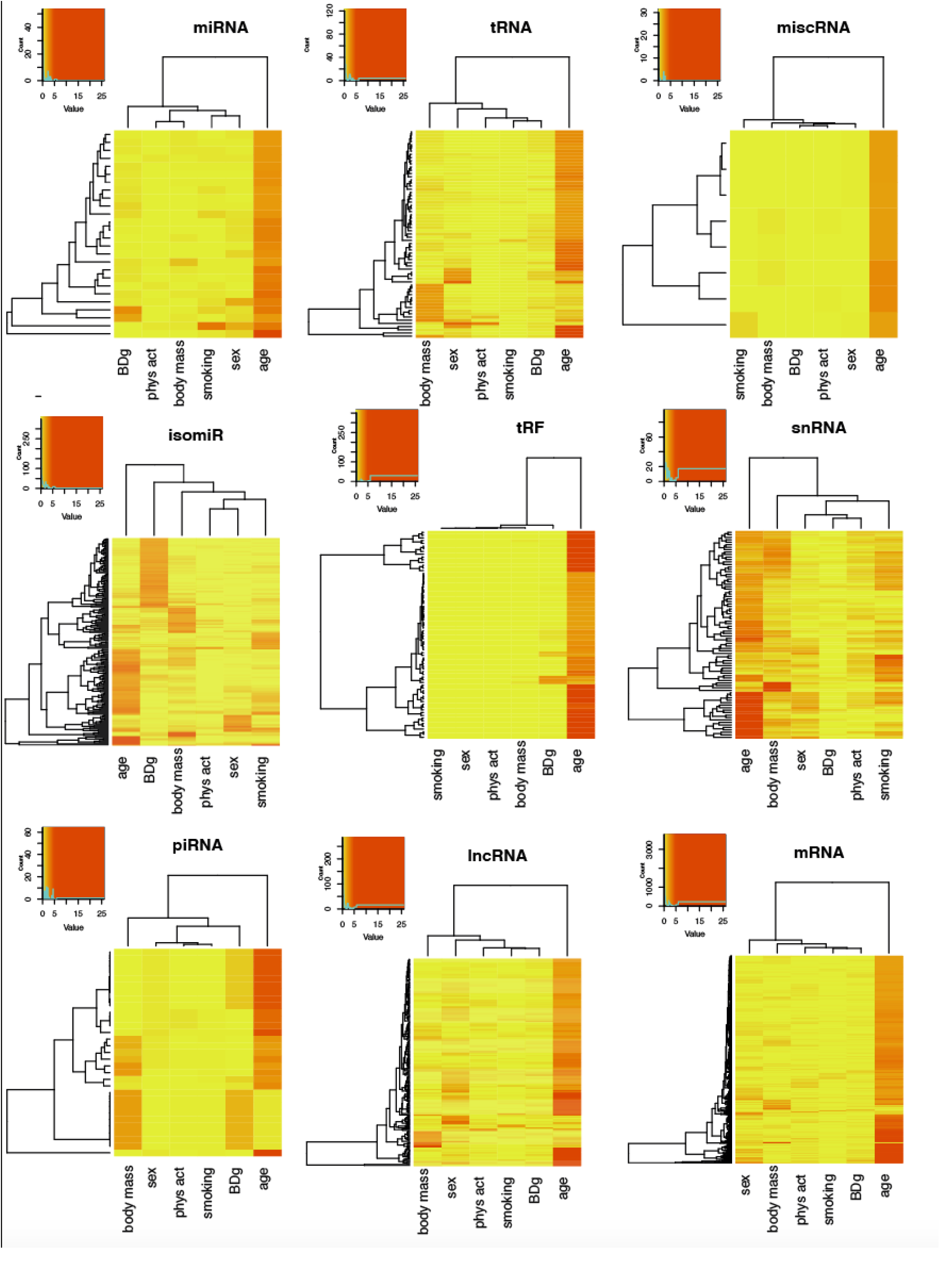
Heatmaps of the hierarchical clustering of -log10 p-values adjusted for multiple testing from the associations between sncRNAs from the classes miRNAs, isomiRs, tRNAs, tRNA fragments, piRNAs, lncRNAs, miscRNAs, snRNAs and mRNA fragments and the attributes blood donor group (BDg), sex, body mass, smoking (current vs never smokers) and physical activity (low vs high activity). sncRNAs are visualized if any of the associations produced p-values <0.01. Colors are yellow to orange for -log10 p-value 0 to 5 and red for -log10 p-values >6.

### Expression differences

The majority of sncRNAs showed log_2_fold transformed differences between 1 and -1 (Figure 3). However, numerous associations with age, and to a lesser extent blood donor groups, showed differences greater than +/-1. Specifically, these larger differences were seen for: miRNAs and blood donor group and age; isomiRs and smoking, age and blood donor group; snRNAs and age; mRNA fragments and age. The majority of miRNAs were upregulated with age while the majority of mRNA fragments were downregulated with age. snRNAs and lncRNAs were also downregulated with age

**Figure 3:**
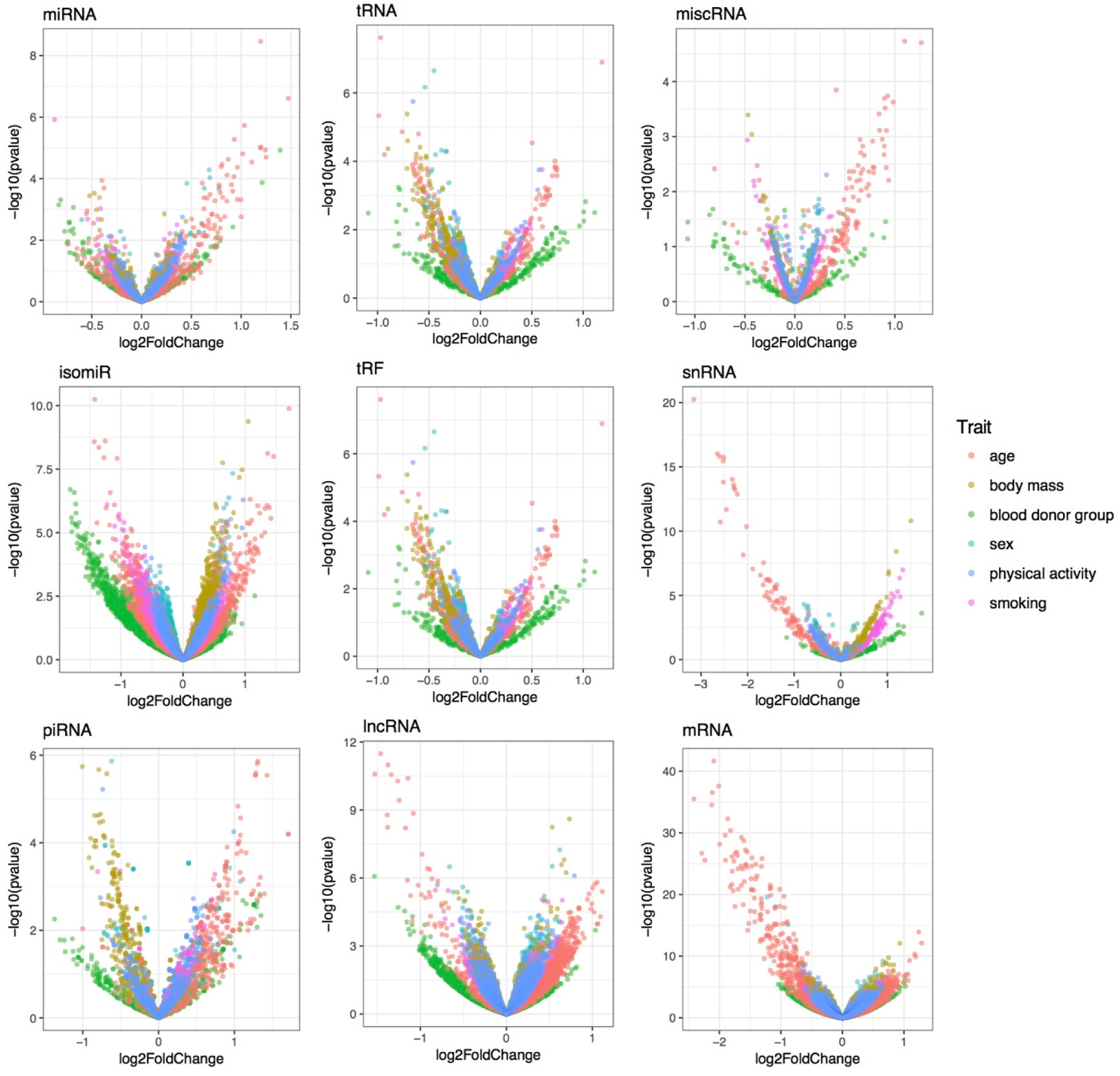
Volcano plots showing differential expression in log_2_fold change on the x-axis and adjusted p-values from the associations in -log10 on the y-axis for miRNAs, isomiRs, tRNAs, tRNA fragments, piRNAs, lncRNAs, miscRNAs, snRNAs and fragments mapping mRNA and the traits; blood donor group (BDg), sex, body mass smoking (current vs never smokers) and physical activity (low vs high activity).

Adjusting for age in the association analyses with sex, body mass, smoking and physical activity showed larger log_2_fold differences compared to unadjusted analyses (Supplementary Figure S2). The changes were striking when compared to the unadjusted analyses (Figure 4), specifically for sncRNA associations with sex. miRNAs and piRNAs were upregulated and lncRNAs and mRNA fragments downregulated in men. Strong smoking differences between current and never smokers on sncRNA expression were also seen for tRNAs, isomiRs, tRNA fragments, snRNAs, lncRNAs and one piRNA. One sex and two body mass associations with mRNAs fragments have log_2_fold differences larger than two and -log10 adjusted p-values >40.

**Figure 4:**
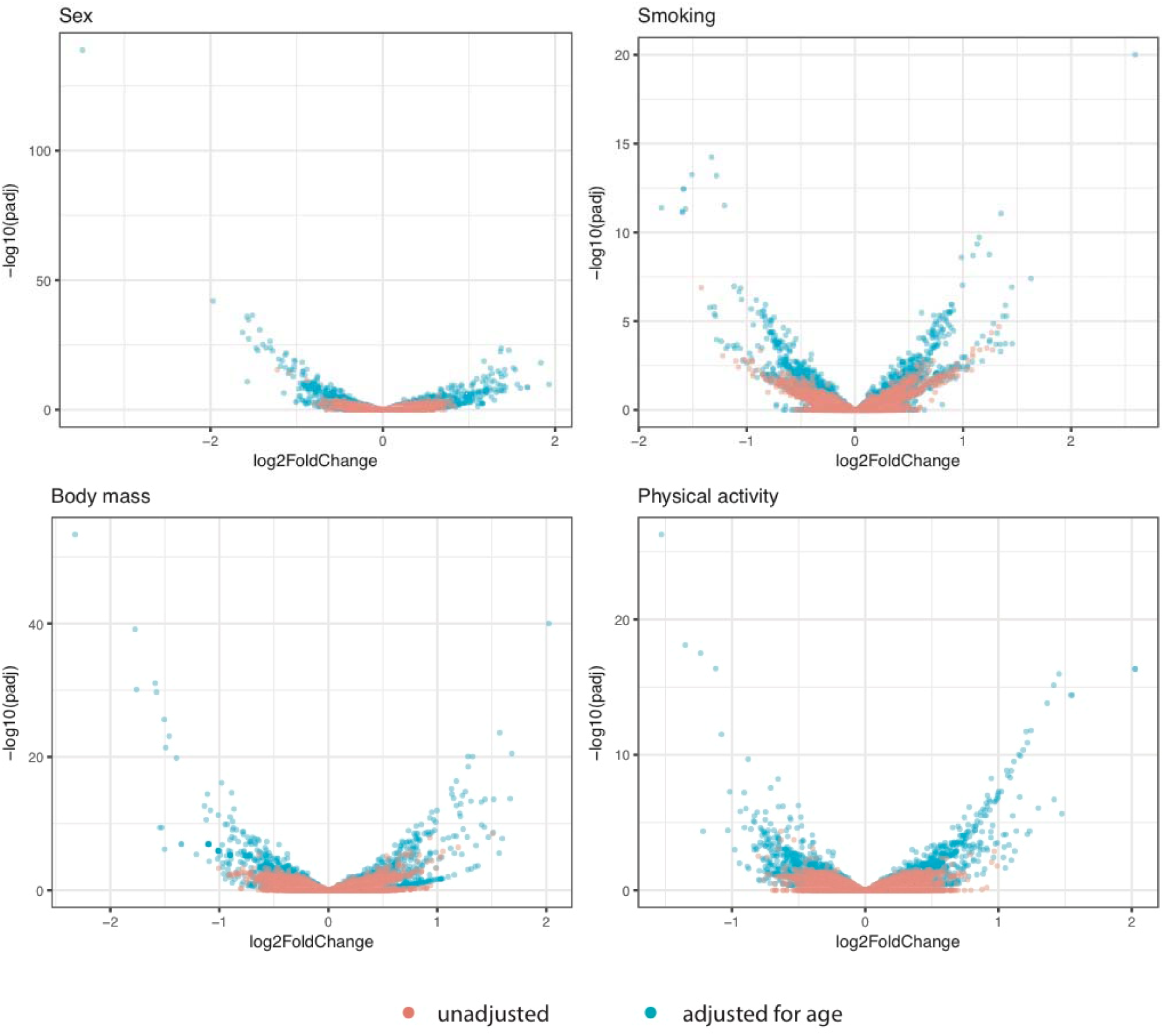
Volcano plots showing differential expression in log_2_fold change on the x-axis and adjusted p-values from the associations in -log10 on the y-axis for all sncRNAs associated to sex, smoking, body mass and physical activity. Associations without age as co-variable are shown in red and associations adjusted for age are shown in blue.

### Co-expression module analyses

Module analyses showed that age and blood donor group are more strongly correlated with co-expression modules than any other trait, followed by physical activity (p-value < 0.01; Supplementary Figure S3). A set of lncRNAs are strongly correlated with sex (Pearson *r*=0.7). A module of 16 tRNA fragments were associated with sex, body mass and physical activity. Smoking is associated with fewer modules than the other traits.

### KEGG Pathway analyses

We performed KEGG pathway analyses for mRNA fragments and miRNA targets (see Materials and Methods). Pathways were enriched for age-, sex- (age adjusted) and smoking-associated (age adjusted) mRNA fragments and miRNA targets (p-value < 0.05; Table 2). We did not detect any significant pathway enrichments for body mass and physical activity. Four out of the top five age-related pathways are involved in carcinogenesis, while all the top five smoking-related pathways have been associated with smoking.

**Table 2:** Pathway enrichment analyses for miRNA targets and mRNA fragments. The top 5 significant enriched pathways for miRNA targets and mRNA fragments for the associations with age, sex adjusted for age and smoking adjusted for age.

| | Age | $-\log_{10}(\text{FDR})$ | Sex, age adjusted | $-\log_{10}(\text{FDR})$ | Smoking, age adjusted | $-\log_{10}(\text{FDR})$ |
| --- | --- | --- | --- | --- | --- | --- |
| 1 | Ras signaling pathway | 1.89 | Axon guidance | 2.45 | Cholinergic synapse | 2.20 |
| 2 | PI3K-Akt signaling pathway | 1.89 | Focal adhesion | 2.45 | Thyroid hormone signaling pathway | 2.20 |
| 3 | Dopaminergic synapse | 1.75 | Endocytosis | 2.16 | Relaxin signaling pathway | 2.02 |
| 4 | MAPK signaling pathway | 1.69 | MAPK signaling pathway | 1.75 | Phosphatidylinositol signaling system | 1.51 |
| 5 | AMPK signaling pathway | 1.69 | Phospholipase D signaling pathway | 1.75 | VEGF signaling pathway | 1.51 |

### Smoking and sncRNA associations

To identify sncRNAs associated with smoking cessation, we assessed differential expression in never vs current smokers relative to never vs former smokers. We identified smoking-related differential expression in isomiRs, piRNA, lncRNA, tRNA and mRNA fragments which persist after smoking cessation. A single piRNA and two tRNAs show persistent expression differences after smoking cessation, while two smoking-associated miRNAs revert to never-smoking levels (Figure 5).

**Figure 5:**
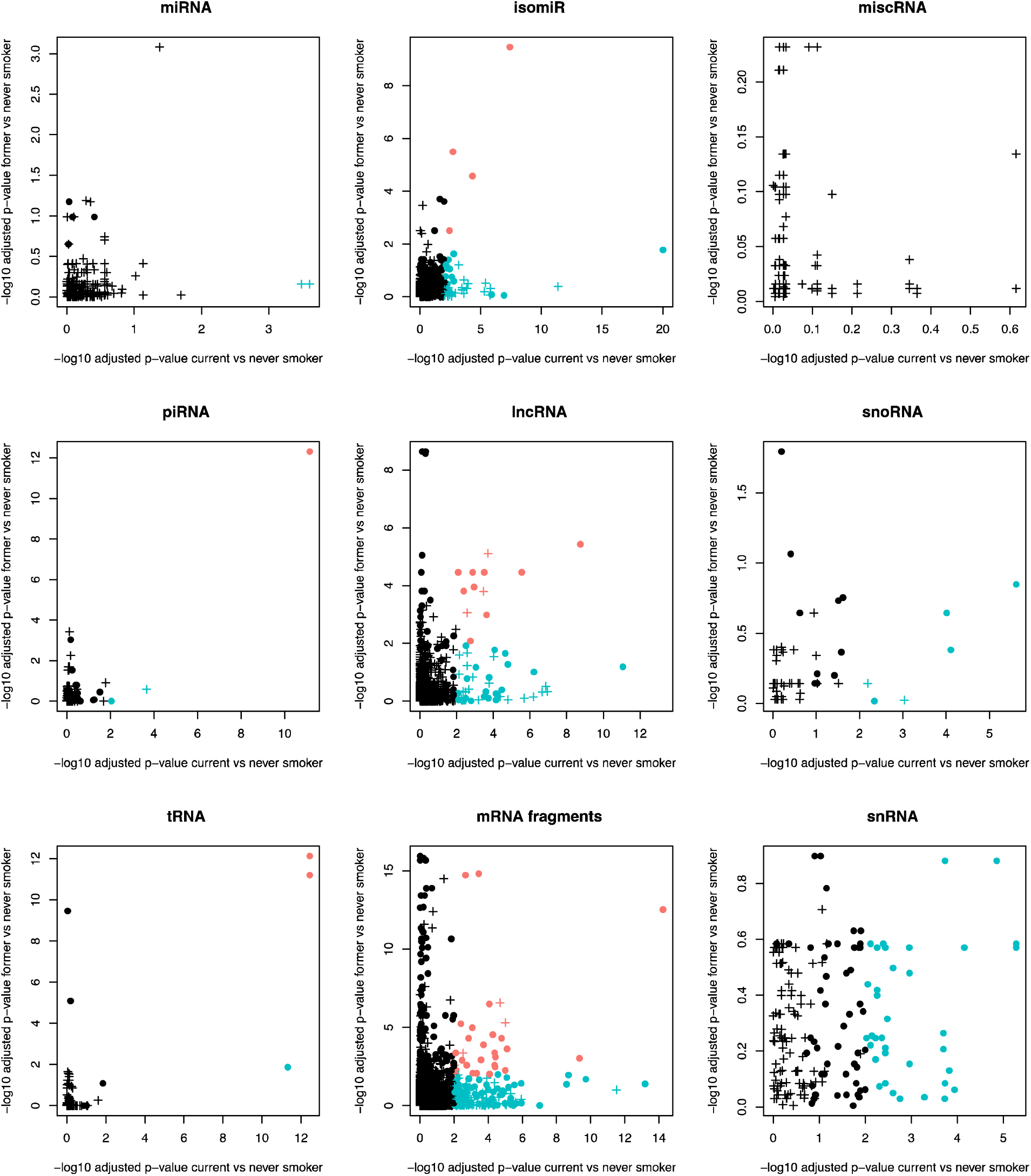
Differential sncRNA expression in never vs current smokers relative to never vs former smokers suggesting smoking related sncRNA expression that persist after smoking cessation (upper right corner) and sncRNA expression that revert to never smoking levels (lower right corner). The -log10 p-value are shown for smoking associations on in current smokers vs never smokers (x-axis) and the former smoker vs never smokers (y-axis). -log10 p-values >2 in both analyses are marked in red, signifying associations both in current and former smokers. -log10 p-values >2 in current, but not in former smokers are marked in blue, signifying associations in current smokers and not in former smokers. Associations with expression differences more than +/-0.5 in both analyses are marked with a cross, all other relationships are marked with a dot. The analyses were done for miRNAs, isomiRs, tRNAs, tRNA fragments, piRNAs, lncRNAs, miscRNAs, snRNAs and mRNA fragments.

The top three smoking-associated miRNAs show a slight increase in expression levels between never, former and current smokers (Figure 6A). This effect became more pronounced in heavy smokers, specifically for individuals smoking more than 20 cigarettes per day for miR-3656 and miR-7704 (Figure 6B).

**Figure 6:**
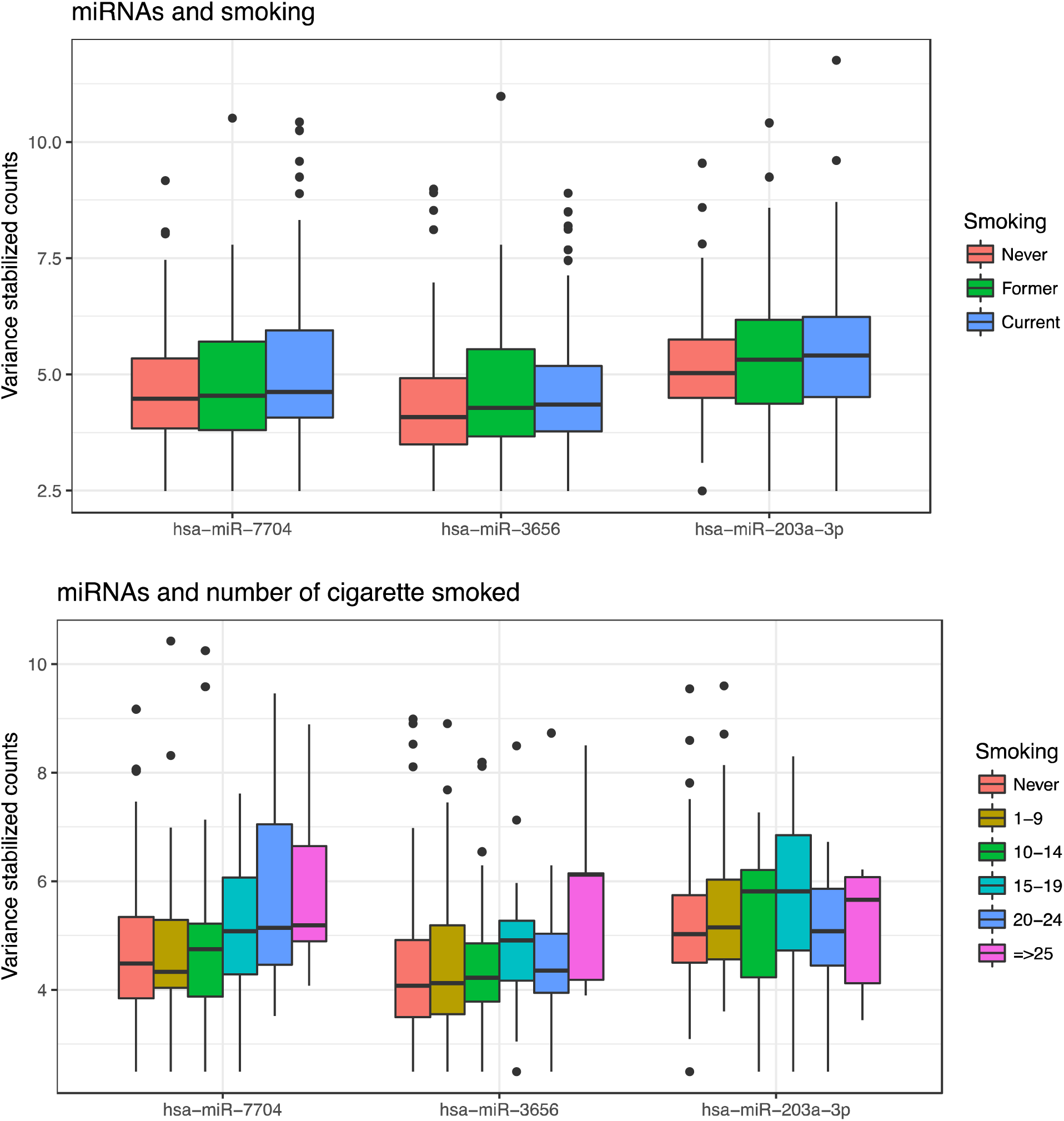
The top 3 smoking associated miRNAs relative to smoking status and dose. A) Boxplot of variance stabilized RNA counts in never former and current smokers of the top three miRNA with lowest p-values. B) Boxplots of variance stabilized RNA counts relative to the number of cigarettes smoked per day.

## Discussion

Expression levels of circulating sncRNAs vary between healthy individuals^46,47^. However, not much is known about which traits influence this variation. Relationships between miRNA expression and age, sex, body mass, smoking and physical activity have been reported^29,34,35,38,41,42^, although most studies have small sample sizes and therefore will be unlikely to detect subtle changes in expression. Furthermore, very little has been reported about these traits in relationship to expression levels of other sncRNA classes.

In this paper, sncRNA association analyses show that ageing is strongly correlated with all sncRNA classes. The age-association was confirmed by analysing sncRNA co-expression modules. Our study is the largest to date showing a strong age effect for all sncRNA classes. The age effect has been consistently reported in previous studies for miRNAs. However, the reported age-associated miRNAs reported in these studies differ presumably due to differences in biological materials, sample processing and sample size.

Age-associated miRNA expression has previously been shown in model organisms^30^, tissues^48^ and blood^29^. In agreement with our results, ageing was reported to be more strongly associated with miRNA expression than sex^29^. miRNA-320b was found to be age-related in both our study and in a large study on whole blood^31^. We found that the age-associated pathways are mostly signalling pathways such as Ras, PI3K-Akt, MAPK and AMPK. This may explain the role of aging in oncogenesis. Ageing is also known to affect dopamine receptors which can explain enrichment of dopaminergic synapse pathway for aging.

The relationship between age and circulating sncRNA expression implies that all sncRNA biomarker studies should take age at sampling into consideration when analysing and interpreting results. Sample groups should be age-matched, stratified by age, or age-adjusted. sncRNAs mediate a number of cellular functions, and age-associated expression changes may implicate these in ageing processes. Changes in blood cell counts with age^49,50^ may explain some of the differential sncRNA expression. The present study provides a valuable data set for studying mechanisms of ageing and age-related diseases such as cancer.

Our data also showed significant associations between sex, body mass, smoking and physical activity and the expression levels of 1240 sncRNAs after adjusting for age. Age is an important effect modifier for these associations since the differences with and without age-adjustment increase significant associations more than 10-fold.

We observed sex-related expression for all sncRNA classes. miRNA expression correlated with sex has previously been shown^23,29^ and in some cases directly or indirectly linked to hormonal regulation^24–26^. piRNAs were initially thought to be specific to germ cells^51,52^, however circulating piRNAs have recently been identified at significant levels^11,53^. Our dataset identified a large number of RNAs mapping to piRNA databases. JSB serum samples were stored at -25°C for up to 40 years^47^, indicating that piRNAs are stable. The cellular origin of the piRNAs is unknown. We observe a difference in expression between males and females for a large fraction of piRNAs, indicating that some of the circulating piRNAs might originate from germ cells. Our data also showed sex-specific differences in lncRNAs and mRNA fragments. For example, one lncRNA co-expression module is highly correlated with sex and includes Y chromosome-derived lncRNA fragments. Based on our findings, matching or adjusting for sex in differential expression studies may be crucial.

Our data show that smoking alters expression levels for all classes of sncRNAs, which had previously only been shown for miRNAs^34,35^. Wang *et al*.^35^ indicated that only a portion of the smoking-related miRNAs revert to a never smoker expression level after smoking cessation. In contrast, our analyses indicate that miRNA expression in former smokers is similar to never smokers. Futhermore, the expression levels of isomiRs, lncRNAs and mRNA fragments, as well as two tRNA and one piRNA, are significantly different in former smokers compared to never smokers, indicating smoking-related expression persists after smoking cessation for these sncRNAs. Similar results have been shown for DNA methylation^54,55^. It is noteworthy that the top three smoking-related miRNAs (hsa-miR-7704, hsa-miR-3655 and hsa-miR-203-3p) showed a clear relationship between smoking-dose and expression levels, and the top five age-adjusted smoking related pathways are involved with smoking. For example, the cholinergic synapse pathway is associated with nicotine addiction^56^ and the relaxin signaling pathway is disrupted by smoking^57^. Our results show that smoking-related pathway RNAs (e.g. mRNAs and miRNA targets) can be identified in serum.

Body size was associated with 411 sncRNAs of which 63% are mRNA fragments. No miRNAs were statistically significant (P<0.001). 208 sncRNAs were associated with physical activity, of which 70% mapped to mRNA fragments. Notably, the overlap between body mass and physical activity related sncRNAs was observed for three tRNAs and 13 tRNA fragments. To our knowledge, no comparable study is available showing physical activity and body mass associations with circulating non-miRNA sncRNAs. Our results indicate that differential expression studies in obesity and exercise should consider studying other sncRNAs in addition to miRNAs.

The serum samples were stored long-term at -25°C. Under these conditions all unstable RNAs have been degraded. We have shown that the total amount of miRNA was affected by the processing of the serum and to a lesser extent by storage time^21^. Also, the number of other sncRNAs decrease with storage^47^. The differential expression analyses between the blood donor groups shown here shed further light on which sncRNAs are affected by storage and processing and further sncRNA studies using JSB will take blood donor group into consideration.

The primary functions of most RNA classes are known. For example, snRNAs are involved in mRNA splicing, tRNAs decode mRNAs into peptides, snoRNAs carry chemical modifications to mRNA fragments, miRNA regulate post-transcriptional gene expression, piRNAs target and repress the expression of transposable elements and lncRNAs provide epigenetic control of gene expression and promoter-specific gene regulation^58–61^. However, secondary functions are largely unknown and therefore pathways and network approaches for functional analyses are not yet feasible. Another challenge in the interpretation of the results are insufficient accuracy and completeness of the annotation databases. Recognized databases such as miRBase^62^, ENCODE^1^ and piRbase^63^ may include degradation products, misclassifications and mapping errors. Curated databases such as miRgenedb^64^ may improve the interpretability. However, the discovery of sncRNA biomarkers are less affected by poor annotation. piRNAs are particularly difficult and there is a highly probable that the available piRNA databases contain RNAs unrelated to the piRNAs produced by germ cells.

Circulating sncRNAs originate from multiple cell types, and cell type compositional differences might introduce variation or confounding. However, it is not known if all cell types display age-related miRNA expression^65^, and only small expression differences of cell type composition were seen in one of the largest studies to date^31^. In addition, traits such as obesity, low activity and smoking will likely affect RNA expression less than diseases like cancer. Therefore large samples sizes are needed to discover signal over noise.

The main strength of our study is the large sample size. 526 donors included in the study provided sufficient statistical power to detect small differences in expression. Linkage to a complete cancer registry ensures that all donors were free from cancer at least 10 years after sample donation, removing the effects of potential cancer progression on sncRNA expression. Harmonized and quality-assured smoking, body mass and physical activity data improves the accuracy of the measured traits^45^. High sequence read-depth (on average 18 mill reads per sample) serum RNAseq data targeting RNAs between 17 and 47 nucleotides in length enables comprehensive assessment of all main RNA classes.

The primary limitation of the study is the long-term storage of the samples and the effect it might have on RNA quality. Although the advantage of long-term storage is long follow-up time for the disease outcome. The expression differences from storage and sample handling may affect the associations, however, the effects found in previous studies were minor^21,47^. Common with all sncRNA studies, problems with annotation and the lack of functional information makes interpreting the findings challenging. Although the data has unprecedented sample size, the moderate-high activity group and individuals less than 40 years old are represented by fewer than 100 individuals. Associations were calculated from variable samples sizes, due to missing data. This might to some extent reduce comparability between trait associations.

In conclusion, our study showed that sncRNA expression levels in serum are strongly age-dependent, and therefore age should be taken into account in studies of circulating sncRNA expression. sncRNA expression also differed between sexes, and this difference may reflect key biological differences, such as germ cell specificity of piRNAs. Some of the expression signatures are also influenced by body mass, smoking, physical activity and sample processing. The relationships between traits and sncRNA expression levels are of key importance in all sncRNA biomarker research and should be accounted for in the study design and analyses of data.

## Materials and Methods

### Study design

The Janus Serum Bank (JSB) is a population-based cancer research biobank containing prediagnostic biospecimens from 318 628 Norwegians^44^. We identified 550 JSB donors that were cancer free at least 10 years after sample donation using data from the Cancer Registry of Norway. Information on age at donation, processing of samples according to blood donor groups (BDg), sex, body mass, smoking and physical activity were available for the donors (Figure 1A).

Inclusion criteria for each trait were a high-quality sncRNA profile (see filtering criteria in the bioinformatics section) and available trait information. JSB has prospectively collected serum samples between 1972 and 2004. The collection procedure and serum processing have varied throughout the collection period. 156 samples included were red cross blood donors (RCBD) and 370 were donors participating in health examinations (HEBD), in total 526. The samples were grouped according to sample collection period and processing (Grp1:HEBD from 1972-1978, iodoactetate added, Grp2: HEBD from 1979-1986, Grp3: HEBD from 1987-2004 collected in separating gel tubes, Grp4A: RCBD from 1973-1979, Grp4B: RCBD from 1973-1979, lyophilized, Grp5: RCBD from 1980-1990 and Grp6: RCBD from 1997-2004)^21^ (Figure 1B). The number of samples in each blood donor (BD) group from 1 to 6 were 33, 143, 194, 62, 66 and 28, respectively. 171 women and 288 men (Figure 1B) with a mean age of 50 at donation were included. Age at donation was categorized into less than 40, between 40 and 60 and above 60 (Figure 1B). Data from the health examination studies were available for the HEBD donors, including information on smoking habits, body mass index (BMI) and physical activity (Figure 1B)^45^. The donors with smoking information were categorized into current (N=107), former (N=85) and never (N=111) smokers. The number of cigarettes per day in current smokers were categorized into 1-9 (N=36), 10-14 (N=33), 15-19 (N=22), 20-24 (N=19) or >=25 (N=5). There were 152 normal weight, 4 underweight, 156 overweight and 39 obese donors using WHO standards. Analyses were done contrasting normal weight vs overweight and obese combined. The donors characterized their physical activity to be inactive (N=37), low (N=177), moderate (N=70) or hard (N=9). The inactive and low were compared to moderate and hard. (Figure 1B).

This study was approved by the regional committees for medical and health research ethics, Oslo, Norway (2016/1290).

### RNA isolation and sequencing protocols

RNA was extracted from 2 x 200 µl serum using phenol-chloroform separation and the miRNeasy Serum/Plasma kit (Cat. no 1071073, Qiagen) on a QIAcube (Qiagen). Glycogen (Cat. no AM9510, Invitrogen) was used as carrier during the RNA extraction step. The eluate was concentrated using Ampure beads XP (Agencourt). Small RNAseq was performed using NEBNext^®^ Small RNA Library Prep Set for Illumina (Cat. No E7300, New England Biolabs Inc.) with a cut size on the pippin prep (Cat. No CSD3010, Sage Science) covering RNA molecules from 17 to 47 nucleotides. Sequencing libraries were indexed and 12 samples were sequenced per lane on a HiSeq 2500 (Illumina) to an average depth of 18 million reads per sample.

### Bioinformatics analyses

The RNAseq reads were initially trimmed for adapters using AdapterRemoval (v2.1.7)^66^. We then mapped the collapsed reads (generated by FASTX v0.14) to the human genome (hg38) using Bowtie2 (10 alignments per read were allowed). We compiled a comprehensive annotation set from miRBase (v21)^62^ for miRNAs, pirBAse (v1.0) for piRNAs^63^, GENCODE (v26)^1^ for other RNAs and tRNAs. We used SeqBuster (v3.1)^67^ to get isomiR and miRNA profiles. To count the mapped reads, HTSeq (v0.7.2)^68^ was used. The candidate tRNA fragments (tRFs) were selected from the reads mapped to tRNA annotations. For biomarker purposes, we excluded RNAs with fewer than 10 reads in more than 20% of the samples (Figure 1C medium stringent filtering). To show how filtering influenced the number of RNAs we produced tables with stringent and less stringent filtering cut-offs. Stringent filtering excluded RNAs with less than 10 reads in more than 50% of the samples. Less stringent filtering excluded RNAs with less than 1 read in more than 20% of samples (Figure 1C).

### Statistics

Differential gene expression analyses based on the negative binomial distribution and Wald significance tests were performed for each trait using the R package DESeq2 version 1.14.1^69^. All traits were categorical. The analyses were performed with and without adjustment for age at donation. P-values after adjusting for multiple testing, using DESeq2 default adjustments, were reported^69^. Heatmaps of trait-associated RNAs were created using the heatmap.2 function in the gplots package. sncRNAs where any of the traits had adjusted p-value < 0.001 for analyses not adjusted for age at donation and p-values < 0.01 for analyses adjusted for age at donation are shown. We performed variance stabilizing transformation (VST) from the fitted dispersion-mean relations and then transformed the normalized count data using the function varianceStabilizingTransformation. Variance stabilized normalized counts were extracted and in-depth analyses of the top 3 strongest associations for smoking, body mass and physical activity were performed. For this, current, former, never smokers and number of cigarettes per day were investigated. Body size was categorized according to WHO standardized cutoffs and physical activity were analysed according to the levels inactive, low, moderate and high. We performed KEGG pathway^70^ analysis on differentially expressed mRNA fragments and miRNA targets. The analysis was performed using R function *kegga* from the *limma* package. The miRNA targets were extracted from miRDB (v5.0) predictions^71^ (score cut off > 60).

### Co-expression module analysis

We used the weighted correlation network analysis (WGCNA) R package (v1.61)^72^ to determine co-expression modules among serum RNAs. The samples that have any missing values among their traits were filtered out and the remaining samples were utilized for co-expression module identification. The identified modules (min. module size is 10) were mapped to the sample traits to find significantly (p-values < 0.01) correlated associations between the modules and traits. The effect sizes were measured using Pearson correlation coefficients.

## Data availability

Sequence data have been deposited at the European Genome-phenome Archive (EGA), which is hosted by the EBI, under accession number EGASxxxxxxx, with restricted access. Custom scripts are available from the corresponding author upon request.

## Acknowledgements

This study was funded by The Norwegian Research Council’s Programme ‘Human Biobanks and Health Data (project nr 229621/H10 and 248791/H10). We would like to acknowledge Cecilie Bucher-Johannessen, Marianne Lauritzen, Magnus Leithaug for performing lab and coordination tasks and Ronnie Babigumira and Jan Ivar Martinsen for data management. The sequencing service was provided by the Norwegian Sequencing Centre (www.sequencing.uio.no), a national technology platform hosted by Oslo University Hospital and the University of Oslo supported by the Research Council of Norway and the Southeastern Regional Health Authority.

## Author contributions

TBR and HL designed the study, TBR and SUU performed the analyses of the data, TBR, RL and HL wrote the draft of the paper, TBR, SUU, AK, EM, GU, ST, RL and HL discussed the results and contributed to the writing of the final manuscript.

